# Imagined speech can be decoded from low- and cross-frequency features in perceptual space

**DOI:** 10.1101/2021.01.26.428315

**Authors:** Timothée Proix, Jaime Delgado Saa, Andy Christen, Stephanie Martin, Brian N. Pasley, Robert T. Knight, Xing Tian, David Poeppel, Werner K. Doyle, Orrin Devinsky, Luc H. Arnal, Pierre Mégevand, Anne-Lise Giraud

**Author notes:** Authors contributed equally to this work (Timothée Proix, Jaime Delgado Saa). Senior authors contributed equally to this work (Luc H. Arnal, Pierre Mégevand, Anne-Lise Giraud).

## Abstract

Reconstructing intended speech from neural activity using brain-computer interfaces (BCIs) holds great promises for people with severe speech production deficits. While decoding *overt* speech has progressed, decoding *imagined* speech have met limited success, mainly because the associated neural signals are weak and variable hence difficult to decode by learning algorithms. Using three electrocorticography datasets totalizing 1444 electrodes from 13 patients who performed overt and imagined speech production tasks, and based on recent theories of speech neural processing, we extracted consistent and specific neural features usable for future BCIs, and assessed their performance to discriminate speech items in articulatory, phonetic, vocalic, and semantic representation spaces. While high-frequency activity provided the best signal for overt speech, both low- and higher-frequency power and local cross-frequency contributed to successful imagined speech decoding, in particular in phonetic and vocalic, i.e. perceptual, spaces. These findings demonstrate that low-frequency power and cross-frequency dynamics contain key information for imagined speech decoding, and that exploring perceptual spaces offers a promising avenue for future imagined speech BCIs.

## Introduction

Cerebral lesions and motor neuron disease can lead to speech production deficits, or even to a complete inability to speak. For the most severely affected patients, decoding speech intentions directly from neural activity with a BCI is a promising hope. The goal is to teach learning algorithms to classify and decode neural signals from imagined speech, e.g. syllables, words, and to provide feedback to the patient so that the algorithm and the patient adapt to each other. This strategy parallels what is being done in the motor domain to help paralyzed people control e.g. a robotic arm (Hochberg et al., 2012). One approach to decode imagined speech is to train algorithms on articulatory motor commands produced by the brain during overt or silently articulated speech, hoping that the learned features could ultimately be transferred to patients who are unable to speak (Anumanchipalli et al., 2019; Livezey et al., 2019; Makin et al., 2020). Although potentially interesting, this hypothesis is limited in scope as it would only work for those cases where language and motor commands are preserved, such as in motor neuron disease, i.e. in a minority of the patients with severe speech production deficits (Guenther et al., 2009; Wilson et al., 2020). If, as in most cases of post-stroke aphasia, the cortical language network is injured, other decoding strategies must be envisaged, for instance using neural signals from the remaining intact brain regions that encode speech, e.g. regions involved in perceptual or lexical speech representations. Exploring these alternative hypotheses require to work directly from imagined speech neural signals, even though they are notably difficult to decode, because of their high spatial and temporal variability, their low signal-to-noise ratio, and the lack of behavioral outputs. To advance imagined speech decoding, two key points must be clarified: (i) what brain region(s) and associated representation spaces offer the best decoding potential, and (ii) what neural features (e.g. signal frequency, cross-frequency or -regional interactions) are most informative within those spaces.

Until now, imagined speech decoding with non-invasive techniques, i.e. surface EEG/MEG, has only led to poor results (Bocquelet et al., 2016). The most promising approach is based on electrocorticographic (ECoG) signals, which, so far, are only recorded in patients with refractory epilepsy undergoing presurgical evaluation. During the experiment, patients are typically asked to speak aloud or imagine speaking or hearing, and ECoG signals are recorded simultaneously. In the *overt* condition, the recorded speech acoustics is used to inform the learning algorithms about the timing of speech production in the brain. The main state-of-the-art feature used for overt speech decoding is the broadband high-frequency activity (BHA) (Leszczyński et al., 2020; Rich and Wallis, 2017). When sampled from the premotor and motor articulatory cortex (Chartier et al., 2018; Ray and Maunsell, 2011; Steinschneider et al., 2008), this feature permits reasonable decoding performance. However, even though patients have an intact language and speech production system (Martin et al., 2016, 2014), this feature is less efficient when speech is imagined. Alternative features or feature combinations are hence needed to advance from decoding overt speech to the more clinically relevant step of decoding imagined speech.

The feature space being potentially unlimited, it is essential for future treatment of aphasia to reduce the amount of exploited features to the most promising ones, as for prophylactic reasons intracortical sampling will have to remain as restricted as possible. Existing speech and language theories, in particular, theories of imagined speech production, can help us target the best speech representation level(s) and associated brain regions. While the motor hypothesis posits that imagined speech is essentially an attenuated version of overt speech with a well specified articulatory plan (much like imagined and actual finger movements share similar spatial organization of neural activity), the abstraction hypothesis proposes that it arises from higher-level linguistic representations that can be evoked without an explicit plan (Cooney et al., 2018; Indefrey and Levelt, 2004; Mackay et al., 1992; Miller et al., 2010; Oppenheim and Dell, 2010; Wheeldon and Levelt, 1995). Between these two accounts, the flexible abstraction theory assumes that the main representation level of imagined speech is phonemic, even though subjects can retain control on the contribution of sensory and motor components (Oppenheim and Dell, 2010; Pickering and Garrod, 2013; Scott et al., 2013; Tian, 2010). In this case, neural activity is shaped by the way each individual imagines speech (Perrone-Bertolotti et al., 2014). An important argument for the flexible abstraction hypothesis is that silently articulated speech exhibits the phonemic similarity effect, whereas imagined speech without explicit mouthing does not (Oppenheim and Dell, 2010). Altogether these theories suggest that semantic and perceptual spaces deserve as much attention as the articulatory dimension in imagined speech decoding.

Other current theories of speech processing (Giraud and Poeppel, 2012) may provide important complementary information to identify the best neural features to exploit within those spaces. These theories suggest that that other frequency features than BHA are critical to speech neural processing and encoding (Giraud and Poeppel, 2012). Slower frequencies, in particular the low-gamma and theta bands could underpin phoneme- and syllable-scale processes that are essential for both speech perception and production, such as the concatenation of segment-level information (phoneme-scale) within syllable timeframes. This hierarchical embedding could be operated by nested theta/low-gamma and theta/BHA phase-amplitude cross-frequency coupling (CFC) both in speech perception and production (Giraud, 2020; Giraud and Poeppel, 2012; Gross et al., 2013; Hovsepyan et al., 2020; Marchesotti et al., 2020). The low-beta range could also contribute to speech encoding as it is implicated in top-down control during language tasks (Lewis and Bastiaansen, 2015; Pefkou et al., 2017). In coordination with other rhythms, such as the low-gamma band, it participates in the coordination of bottom-up and top-down information flows (Bastos et al., 2020; Fontolan et al., 2014; Rimmele et al., 2018).

These frequency specific neural signals could be of particular importance for intended speech decoding, as focal articulatory signals indexed by BHA are expected to be weaker during imagined speech.

In this study, we set out to delineate the range of representation level(s) and neural features that could potentially be usable in imagined speech decoding BCIs. Rather than adopting a purely neuroengineering perspective involving large datasets and automatized feature selection procedures, we used a hypothesis-driven approach assuming a role of low-frequency neural oscillations and their cross-frequency coupling in speech processing, within both perceptual and motor representation spaces.

## Results

Imagined speech experiments were carried out in three groups of participants implanted with ECoG electrodes (4, 4, and 5 participants with 509, 349, and 586 ECoG electrodes for studies 1, 2, and 3 respectively, Fig. 1). Each group performed a distinct task, but all studies involved repeating out loud (overt speech) and imagining saying or hearing (imagined speech) words or syllables, depending on the study (see Methods).

**Figure 1:**
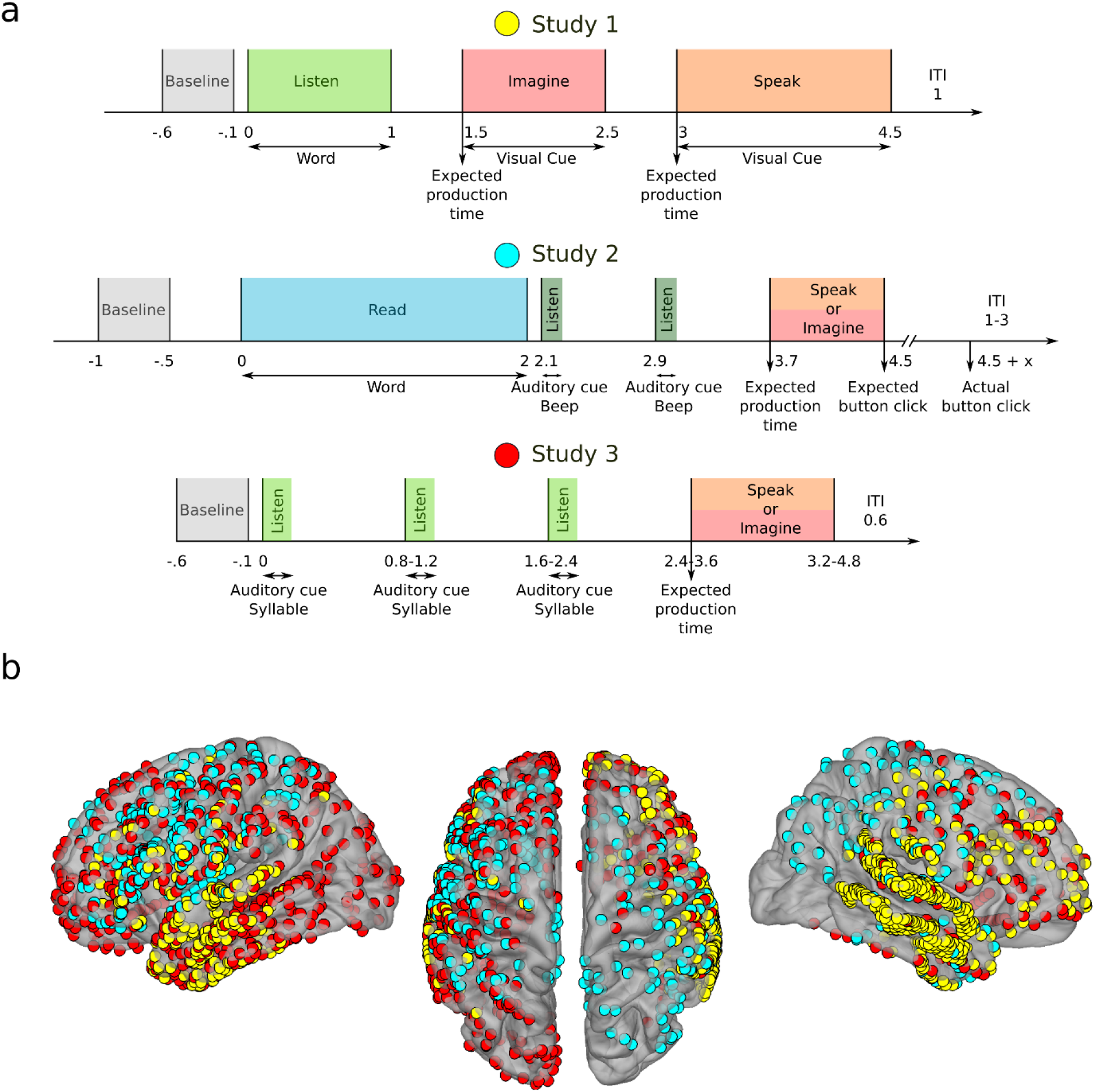
Experimental studies and electrode coverage. (**a**) Study 1 (top row): After a baseline (0.5 s, grey), participants listened to one of six individual words (1 s, light green). A visual cue then appeared on the screen, during which participants were asked to imagine hearing again the same word (1 s, red). Then, a second visual cue appeared, during which participants were asked to repeat the same word (1.5 s, orange). Study 2 (middle row): After a baseline (0.5 s, gray), participants read one of twelve words (2 s, blue). Participants were then asked to imagine saying (red) or to say out loud (orange) this word following the rhythm triggered by two rhythmic auditory cues (dark green). Finally, they would click a button, still following the rhythm, to conclude the trial. Study 3 (bottom row): After a baseline (0.5 s, gray), participants listened to three rhythmic auditory repetitions of the same syllable (light green) with different rhythms speeds, after which they were asked to imagine saying (red) or to say out loud this syllable (orange). (**b**) ECoG electrode coverage across all participants. Different colors correspond to the three studies.

### Speech item discrimination from power spectrum and phase-amplitude cross-frequency coupling

We first quantified power spectrum changes during overt or imagined speech compared to baseline for four frequency bands: theta (θ, 4-8 Hz), low-beta (lβ, 12-18 Hz), low-gamma (lγ, 25-35 Hz), and BHA (80-150 Hz). Overall, spatial patterns of power spectrum changes for overt and imagined speech were comparable, but not identical. Furthermore, power changes for imagined speech were less pronounced than those for overt speech, with fewer cortical sites showing significant changes. We found power increases in the BHA for both overt and imagined speech in the sensory and motor regions (Fig. 2), and power decrease in the beta band over the same regions. A smaller power decrease was also found over the same regions for theta and low-gamma band (Supp. Fig. 1). The most striking difference between overt and imagined spatial patterns was that BHA in superior temporal cortex increased during overt speech whereas it decreased during imagined speech, a finding that presumably reflects the absence of auditory feedback in the imagined situation. The differences in power spectrum changes between overt and imagined speech were sufficient to accurately classify which task the participants were engaged in (Supp. Fig. 2).

**Figure 2:**
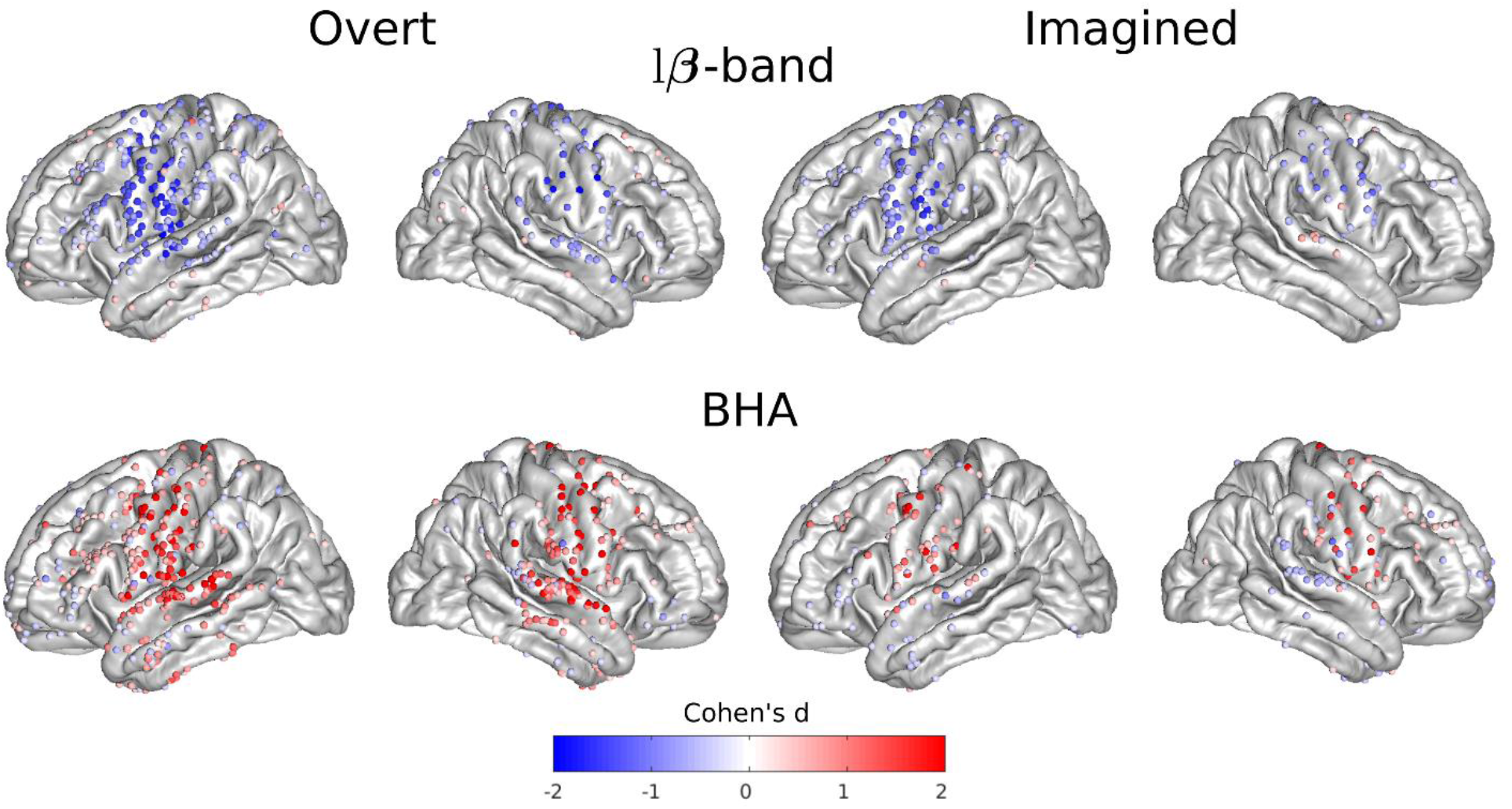
Spatial organization of power spectrum deviations from baseline elicited by overt and imagined speech. Effect sizes (Cohen’s d) for significant cortical sites across all participants and studies during overt and imagined speech compared to baseline (t-tests, FDR-corrected, target threshold *α* = 0.05).

We then quantified phase-amplitude CFC for each cortical site for overt and imagined speech, using the difference in modulation index between speech and baseline periods, for theta, low-beta, and low-gamma modulating (lower) frequency bands, and beta (β: 12-25 Hz), gamma (γ: 25-50 Hz), and BHA modulated (higher) frequency bands. This difference was expressed as a z-score relative to its distribution under the null hypothesis, generated with surrogate data using permutation testing. The spatial pattern of cortical sites displaying significant CFC was more widespread than that of power changes. Notably, strong phase amplitude CFC was found in the left inferior and right anterior temporal lobe between theta phase and other band amplitudes, both for overt and imagined speech (Fig. 3, see Supp. Fig. 3 for other bands).

**Figure 3:**
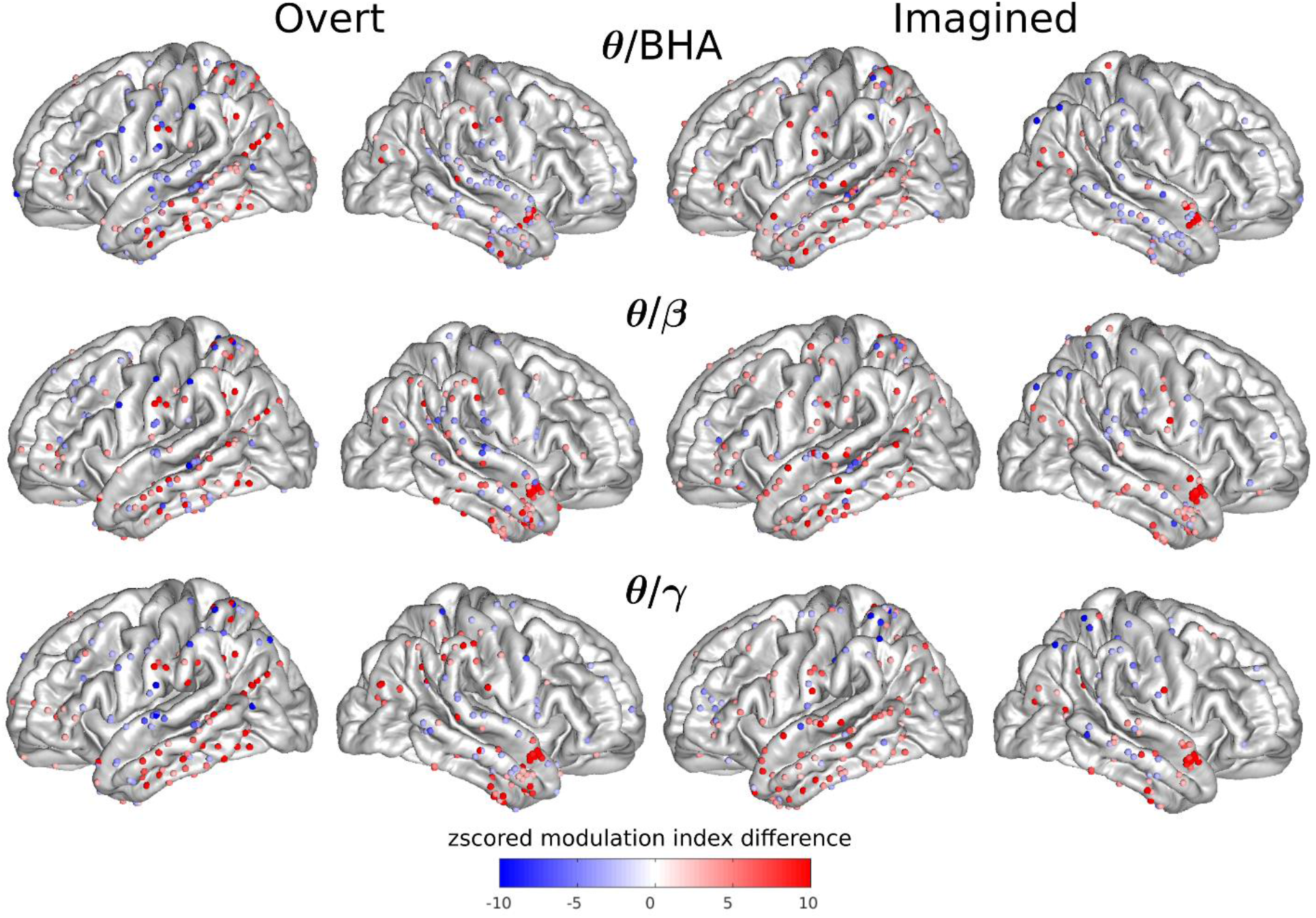
Cross-frequency coupling between the phase of one frequency band and the amplitude of another frequency band for each electrode. Z-scored modulation index difference for significant electrodes across all participants and studies during overt and imagined speech with respect to baseline (permutation tests, FDR-corrected, target threshold *α* = 0.05).

Next, we asked if power spectrum and phase-amplitude CFC changes (hereafter called features) contained information that could be used to discriminate between individual speech words (or syllables in the case of study 3, that we hereafter call speech items). We systematically quantified the correlation between the power spectrum features for all pairs of speech items and their corresponding labels for each cortical site, and averaged the resulting correlation across item pairs. As expected, the BHA showed high correlation values for overt speech, primarily within the sensory-motor and superior temporal cortices of both hemispheres, as well as in the anterior left temporal lobe (Fig. 4). The theta band also showed significant correlations for overt speech in sensory-motor and superior temporal cortex. For imagined speech, however, correlations were more diffuse, in particular for the BHA, with correlation values observed in the left ventral sensory-motor cortex and bilateral superior temporal cortex were lower than for overt speech. Correlations were also observed in the low-beta band in the left superior temporal and the right temporal lobe of the theta and low-beta bands. The same analysis was repeated using phase-amplitude CFC as the discriminant feature (Supp. Fig. 4), showing modest values of correlation in imagined speech.

**Figure 4:**
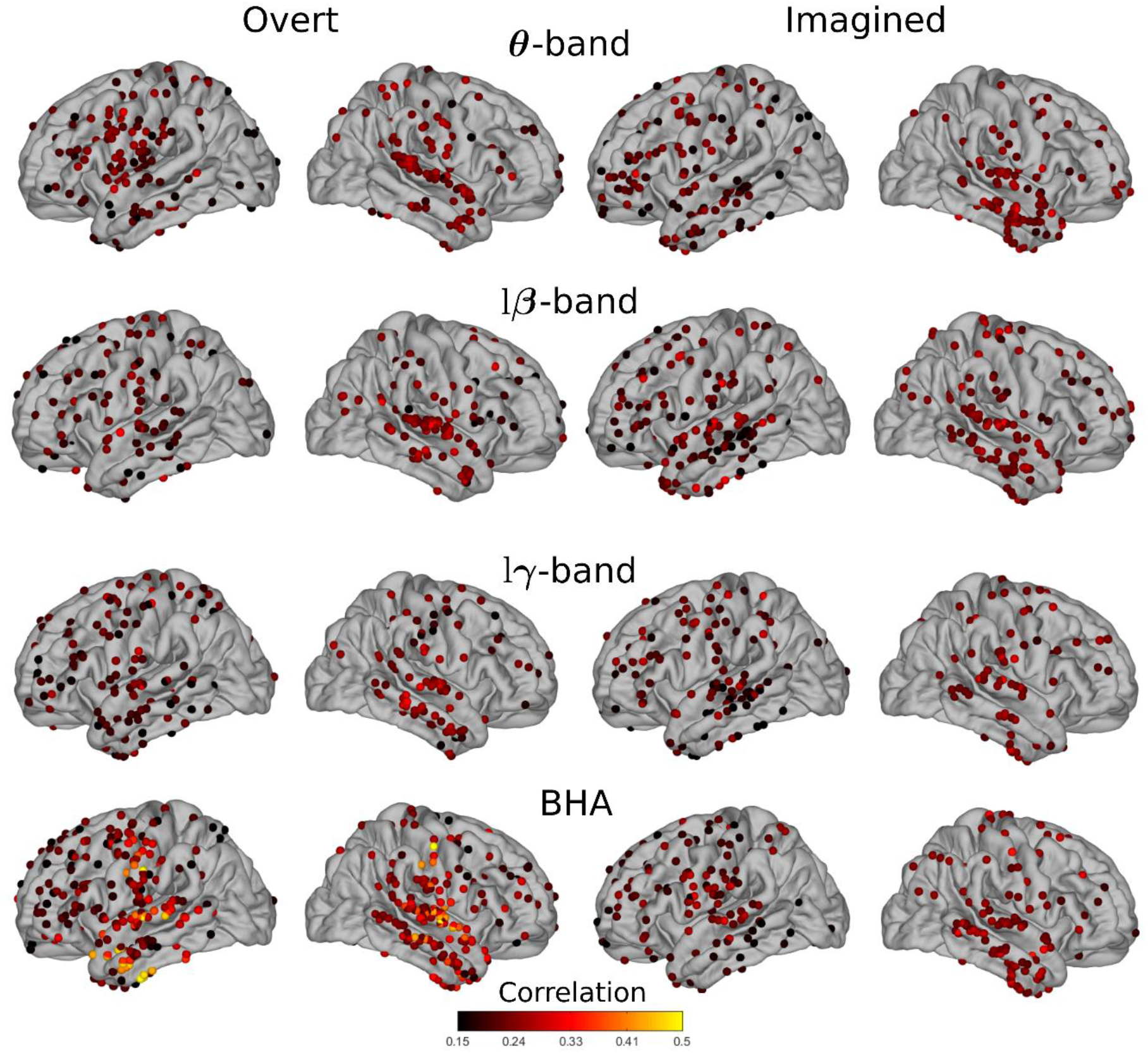
Average correlations between individual speech words and their neural representations. Pairwise correlations between words and power spectrum features averaged across all word pairs for overt and imagined speech for significant electrodes (permutation tests, p<0.05, not corrected for multiple comparison).

### Different articulatory, phonetic and vocalic organization between overt and imagined speech

Based on these initial results, we concluded that the dynamics and neural organization differed for overt and imagined speech production. We therefore asked whether the various spatio-temporal organizations of neural activity during overt speech, i.e. the articulatory organization in ventral sensory-motor cortex (Bouchard et al., 2013; Chartier et al., 2018), the phonetic organization in superior temporal cortex (Mesgarani et al., 2014), the vocalic organization in sensory-motor and superior temporal cortex, and the semantic-syntactic organization in the ventral temporal lobe were conserved during imagined speech. For this, we quantified how well we could discriminate the classes of each speech representation system (i.e. labial, coronal, and dorsal for articulatory representation; fricative, nasal, plosive, and approximant for phonetic representation; low back, low front, high back, high front, and central for vocalic representation; and concrete verb, abstract verb, concrete word, and abstract noun for semantic-syntactic representation [simply called semantic representation hereafter]; see Methods). For each anatomical region of interest (sensory and motor, middle and inferior temporal, superior temporal, and inferior frontal cortices), we built a high-dimensional feature space for which each axis corresponds to one electrode feature. The dimensionality of this feature space was first reduced with PCA. The Fisher distance (which quantifies features separation) was then computed between each pair of speech items across principal components. As all items were made of one or a sequence of phonemes, and thus belonged to at least one group for each representation, the resulting distance could be attributed to the group(s) that were represented in only one of the two words, i.e. to the discriminant one. For instance, the feature distance between the articulatory representations of “python” ([paɪθən], which includes only labial and coronal phonemes) and “cowboys” ([kaʊbɔɪz], which includes only dorsal, labial, and coronal phonemes), was assigned to the dorsal group, as it is the only discriminant one.

For overt speech, as expected, high Fisher distance values were found using power of the BHA in sensory-motor cortex and in the temporal lobe (Fig. 5, see Supp. Fig. 5 for each group separately). During imagined speech, however, the BHA was associated with much lower Fisher distances. In fact, lower frequency bands (theta, low-beta, low-gamma) displayed similar or even higher values in left and right hemispheres for phonetic, vocalic, and semantic representations.

**Figure 5:**
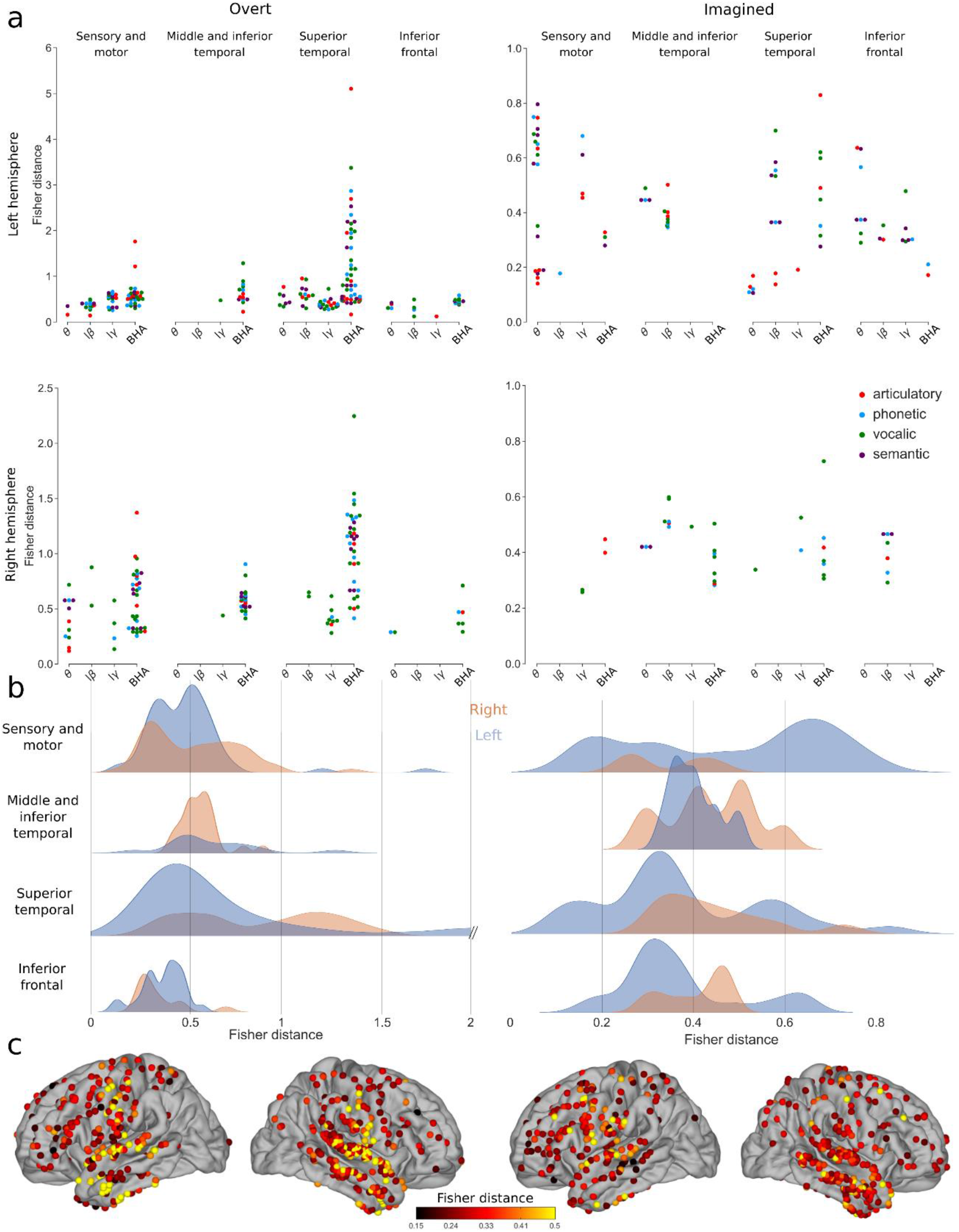
Discriminability between different representations using power spectrum for overt and imagined speech. (**a**) Significant Fisher distance between articulatory, phonetic, vocalic and semantic representations in different brain regions and frequency bands (permutation tests, FDR-corrected, target threshold *α* = 0.05). Note the different scales between overt and imagined speech. (**b**) Distributions of significant Fisher distance for each brain region across all representations and frequency bands (permutation tests, FDR-corrected, target threshold *α* = 0.05). (**c**) Maximum significant Fisher distance for each electrode across all representations and frequency bands. When several significant Fisher distances exist for the same electrode, the maximum value is shown. Only significant electrodes are shown (permutation test, p<0.05, no FDR correction).

Unlike for power spectrum, the Fisher distances for phase-amplitude CFC were in the same range for overt and imagined speech. In the overt speech condition, the highest values were observed for low-beta/gamma phase-amplitude CFC in left sensory-motor and inferior frontal cortex, as well as low-beta/BHA in the left superior temporal lobe (Fig. 6, see Supp. Fig. 6 for each group separately). During imagined speech, high Fisher distances were obtained mainly for low-beta/BHA phase-amplitude CFC in left sensory-motor cortex and right temporal lobe, and for low-beta/gamma CFC in left inferior frontal cortex.

**Figure 6:**
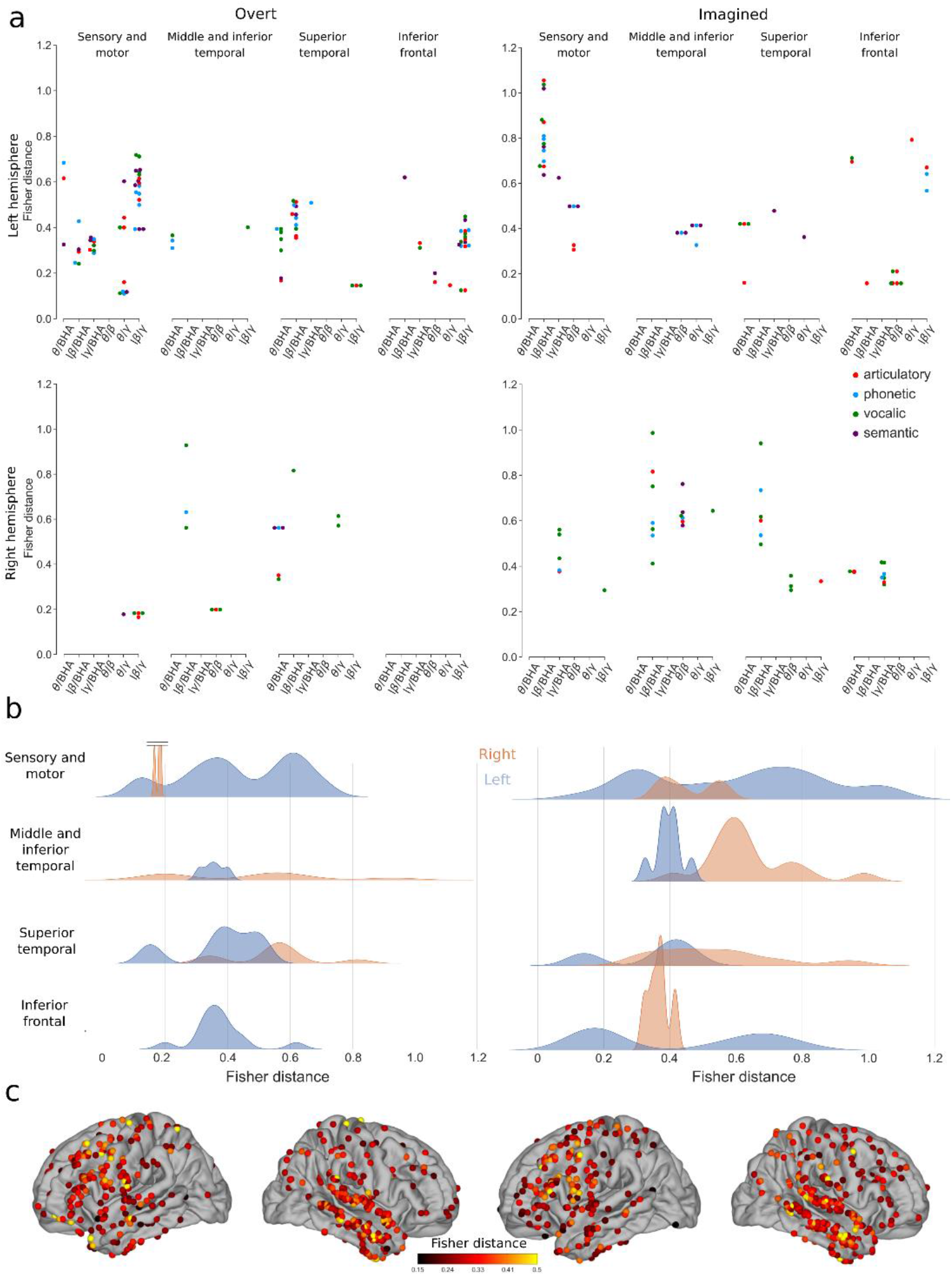
Discriminability between different representations using phase-amplitude CFC changes for overt and imagined speech. (**a**) Significant Fisher distance between articulatory, phonetic, vocalic, and semantic representations in different brain regions and frequency bands (permutation tests, FDR-corrected, target threshold *α* = 0.05). (**b**) Distributions of significant Fisher distance for each brain region across all representations and frequency bands (permutation tests, FDR-corrected, target threshold *α* = 0.05). (**c**) Maximum significant Fisher distance for each electrode across all representations and frequency bands. When several significant Fisher distances exist for the same electrode, the maximum value is shown. Only significant electrodes are shown (permutation test, p<0.05, no FDR correction).

### Decoding imagined speech

Finally, we compared the performance of power spectrum and phase-amplitude CFC for decoding overt and imagined speech (Fig. 7). To simplify the decoding problem and to retain enough trials in each class, we grouped the speech items together to reduce the problem to a binary classification (study 3 was excluded, as it contained only three syllables). New classes were selected by hierarchical clustering of distances between words according to the articulatory, phonetic, and vocalic representations described above (see Methods). Semantic classification was only performed for study 2 by comparing abstract and concrete words.

**Figure 7:**
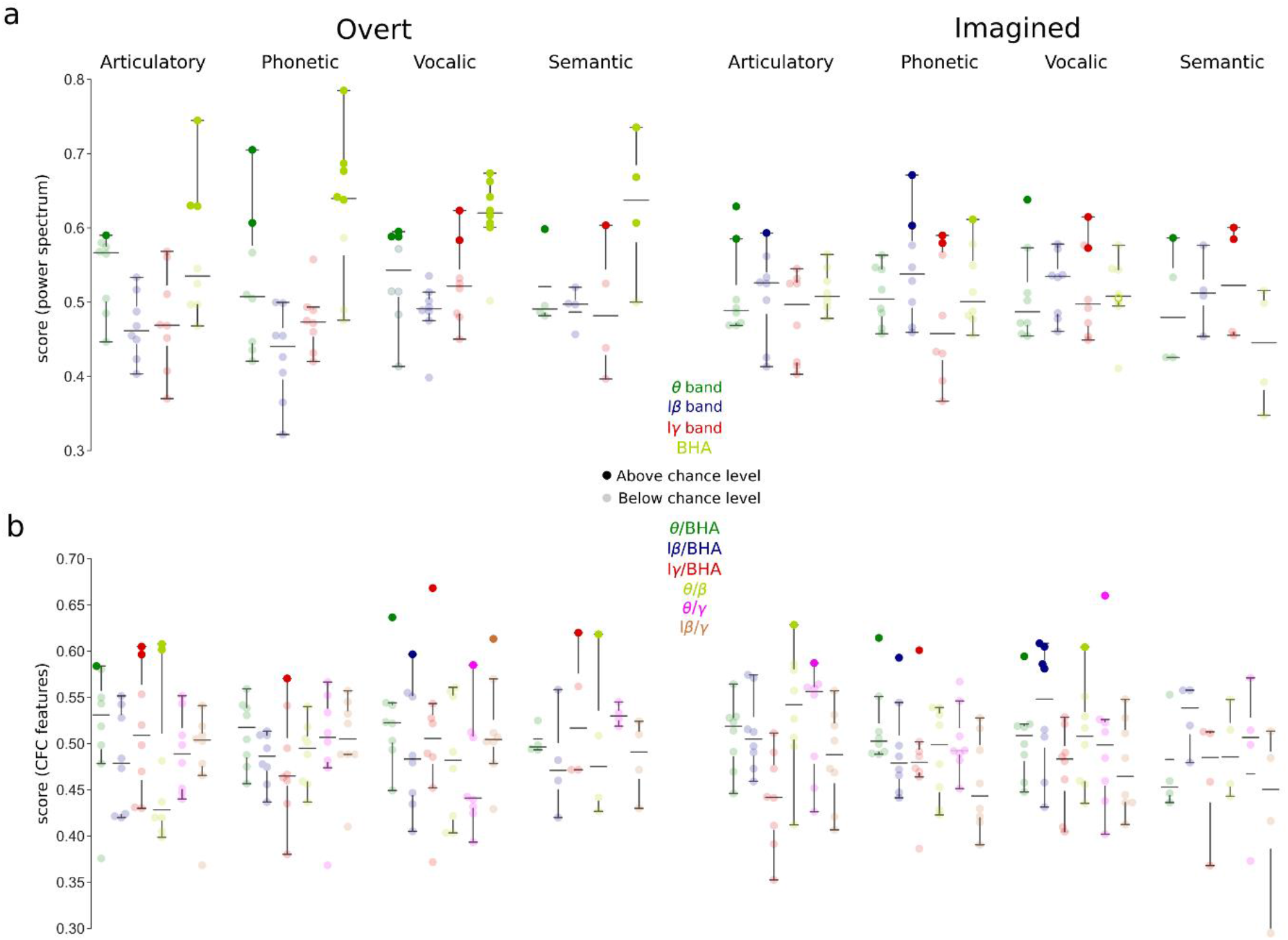
Decoding overt (left) and imagined (right) speech. Opaque (transparent) circles indicate above (below) chance level performance for each participant respectively. For articulatory, phonetic and vocalic decoding, N=8 (studies 1 and 2). For semantic decoding, N=4 (only study 2 had speech items that could be divided into two semantic classes). Boxplot shows the median and interquartile range. Significant levels were obtained for each subject based on the number of trials performed (see Methods) (**a**) Decoding performance using power spectrum features. (**b**) Decoding performance using phase-amplitude CFC features.

For overt speech, good performance could be obtained in 18 participant-representation pairs using power of the BHA, and overall, this frequency band worked better than the others. In imagined speech, however, decoding based on power of the BHA was not better than with other bands. In 13 several participant-representation pairs, classification was as good using e.g., theta or beta power. We also observed that decoding worked better for phonetic and vocalic (i.e. perceptual) representations than for the articulatory one, which supports the flexible abstraction hypothesis of imagined speech. Importantly, the decoding performance for overt speech increased significantly when the trials were realigned using the participant’s voice, suggesting that imagined performance would improve as well if a consistent way of realigning trials could be found (Supplementary note and Supp. Fig. 7).

When using phase-amplitude CFC as a feature, decoding did not perform better for overt (14 participant-representation pairs above chance level) than imagined speech (12 participant-representation pairs above chance level). Participants above chance level were not the same for the different frequency bands and representations. No specific frequency band stood out for overt speech, although the articulatory and vocalic representation worked better. For imagined speech, the low-beta/BHA seems to perform better than other phase-amplitude CFC for imagined speech, confirming the results found in Fig. 6, particularly for the perceptual representations.

## Discussion

In this study, we examined the neural processes underlying the production of overt and imagined speech, in order to identify features that could be used for decoding imagined speech. In particular, we assessed whether these features are similar or different from those that work best for overt speech. To do so we explored not only the articulatory dimension; but also the perceptual (phonetic and vocalic) and semantic representation spaces. We found that overt and imagined speech differ in some crucial aspects of their oscillatory dynamics and functional neuroanatomy. First, while the articulatory representation was well encoded in overt speech, other representations, especially the perceptual one, better reflected imagined speech. Overt and imagined speech both engaged a large part of the left hemispheric language network, with a more prominent involvement of the superior temporal gyrus for overt speech (presumably because of auditory feedback processes). Second, while BHA showed the best performance for overt speech decoding, it conveyed little word- or syllable-specific information during imagined speech. Conversely, neural activity at lower frequencies could be used to decode imagined speech with equivalent or even higher performance than overt speech.

These results suggest that it might prove difficult to successfully transfer the decoding process of brain-computer interfaces trained with overt or even silently articulation speech to imagined speech. BHA representations are poorly specified in primary sensory and motor regions during imagined speech, in accord with the flexible abstraction hypothesis of imagined speech. We also found that the beta-band featured prominently in the neural encoding of imagined speech, both in terms of power and CFC (low-beta/gamma and low-beta/BHA). This finding aligns well with the notion that the beta band plays an important role in endogenous processes, notably in relation with top-down control, in particular in the context of language (Arnal and Giraud, 2012; Bowers et al., 2019; Fontolan et al., 2014; Pefkou et al., 2017). Although repeating a heard or written word engages automatic, almost reflex, neural routines, imagined speech is a more voluntary action requiring enhanced endogenous control from action planning frontal regions (Buschman et al., 2012; Li et al., 2020; Morillon et al., 2019). These results must however be taken with caution as spurious CFC can result from non-linearity, non-stationarity, and power changes across conditions in the signal (Aru et al., 2015; Hyafil, 2015). Even though we carefully selected spectral peaks for the modulating signal to ensure a well-defined phase, and specific bandwidths for the modulated signal, we cannot exclude that significant CFC coupling could theoretically reflect other, non-CFC, changes from baseline to signal. Yet, at the empirical level, that significant and specific decoding performance could be obtained with these features suggests that these frequency features distinguish between speech items, hence contain specific information.

Decoding performance for overt speech increased significantly when trials were aligned on recorded speech onsets (Supp. Fig. 7), which are obviously absent for imagined speech. Previous attempts to align imagined speech directly based on neural data (Martin et al., 2014) met limited success due to the large variability of neural signals across trials and the low signal-to-noise ratio. Although decoding performance would presumably increase if imagined speech onsets and offsets could be detected, we show here that imagined speech decoding is possible using features, such as phase-amplitude CFC, that do not require precise alignment of single-trial data. The absence of behavioral output during imagined speech might even be an advantage, as it definitely prevents the contamination of neural signal recordings by the participant’s voice, a serious problem that was recently discovered. Because the fundamental frequency of the human voice overlaps with the neural BHA, an acousto-electric effect might have artificially inflated the performance in previous overt speech decoding studies (Roussel et al., 2020). To enable a fair comparison of overt and imagined speech in our study, we took care of checking that the three current datasets were free of acoustic contamination. A further technical advantage of silent speech is the absence of movement artefacts. In the three presented studies, the task instructions explicitly stated that participants should not articulate. Using audio/video monitoring, we could confirm that participants did not silently mouth or whisper words, even though it was impossible under our recording conditions to rule out some degree of silent mouthing.

Overall, the current results demonstrate the possibility of obtaining reasonably good decoding performance (>60%) directly from neural activity using electrodes chronically implanted over the cortical surface, and allow us to formulate a number of concrete proposals for the design of future speech BCIs. Using data from three distinct experiments, with similar but not identical task instructions, we could probe the representations of imagined speech at various linguistic levels, namely articulatory, phonological, vocalic and semantic. Despite the typical weakness of imagined speech signals, we reached good decoding performance using lower frequencies and the phonetic representation level. While this is good news for future BCIs, the word level, which was mostly used in this study, is presumably not the optimal currency for an efficient imagined speech decoding strategy based on phonetic representations. A realistic BCI will have to offer decoding based on representation space that can accommodate the size of the average human language repertoire. Likewise, while we showed potential separation in the feature space of syllables, a phoneme decoding strategy would suffer from the combinatorial explosion issue. Using a restricted set of morphemes from which patients could combine to convey the basic needs, could be an interesting first approach. Such a strategy would presumably benefit from the syllable feature space separation shown here. In the future, introducing even more complex, sentence-level stimuli, rather than single words or syllables, could further permit to exploit additional representation levels for imagined speech decoding, such as inference, long-term memory, prosody, semantic mapping, etc. (Gehrig et al., 2019; Huth et al., 2012; Pereira et al., 2018), bringing us closer to ecological and generalizable conditions (Krakauer et al., 2017; Yarkoni, 2019). Each presented stimulus triggers neural activity that might be influenced by word length, frequency, emotional valence, in addition to syntactic and semantic content (Cooney et al., 2018; Pulvermüller, 1999). The richness of these contextual cues could turn out to be an advantage, as it could maximize the separability of speech items, leading to easier decoding, regardless of the representation. In future imagined speech decoding BCIs, specific task instructions will also have to be used to standardize as much as possible imagined speech production. Notably, instructing a participant to “imagine hearing” is expected to induce less residual motion than “imagine speaking”, and to maximally exploit the perceptual representations.

Importantly, our results indicate a large variability in the best decoding features across participants and tasks for imagined speech, suggesting that decoding strategies, i.e. a specific set of spatial and frequency features (anatomical regions, frequency bands, and specific tasks) will have to be adjusted individually in order to build efficient imagined speech BCI systems. In that respect, low frequencies might be more powerful features to decode from spatio-temporally variable signals than BHA, since they tend to be both spatially coherent over larger areas of the cortex, and temporally less constrained. By indexing a more integrated neural activity, they might distinguish better the different imagined speech items. This has practical consequences for the design and placement of future intracranial electrodes. Imagined speech decoding will benefit from a new generation of high-density electrodes that will maximize the amount and quality of the contacts with the cortex. Active multiplexing and graphene-based neural interfaces are two areas of active research in the field (Garcia-Cortadella et al., 2020). With such electrodes and related electronics, on-line signal analysis will be easier, for a more convenient use with BCIs. Off-line analyses such as those we present here are a necessary step to guide us once we will be able to use the novel generation of electrodes in humans and on-line systems. Unlike the robotic arms that are currently being developed for motor restoration, which are optimally controlled by dense sampling of a spatially restricted cortical area (Hochberg et al., 2012), a language BCI system for severe aphasia will require broader coverage of the cortical surface, including the frontal and the temporal lobes, to not only cope with the high physiological intersubject variability of inner speech production, but also with the variable structural damage (cortical, subcortical) that patients may have suffered from. In post-stroke Broca-type aphasia, the efforts to overcome the overt speech planning deficit during imagined speech are expected to implicate a large range of regions of the language network, which will all have to be sampled.

We are just beginning to use machine learning and BCI systems for language restoration, and significant progress can be expected in the coming years, which will lead to unprecedented questions. Among them, the issue of which part exactly of the imagined speech should we let machines decode should trigger careful ethical reflections, which we must conduct ahead of time to prevent abuses and legal loopholes (Rainey et al., 2020). This and other debates, for instance regarding the privacy of neural data, necessitate a multidisciplinary approach that goes beyond the purely technical neuroengineering problem, and pose a challenge that calls for a common effort that we hope scientists will tackle as a community.

## Acknowledgements

This work was funded by the EU FET-BrainCom project, NINDS R3723115, and the Swiss National Science Foundation project grant 163040 to ALG, and by the Swiss National Science Foundation career grant 167836 to PM. The authors thanks Dr. Gerwin Schalk, Dr. Dan Friedman, and Dr. Patricia Dugan for providing access to the datasets used in this work, and Johanna Nicolle for sharing her linguistic expertise.

## Author contributions

S.M., X.T., L.A., P.M., and A.G. designed the experiments. A.C., X.T., and L.A. collected the data. T.P. and J.D. performed the analysis. T.P., L.A., P.M and A.G. drafted the manuscript. All corrected and approved the manuscript.

## Methods

### Participants

Electrocorticographic (ECoG) recordings were obtained in 3 distinct studies from 13 patients (study 1: 4 participants, 4 women, mean age 25.6 years, range 19-33; study 2: 4 participants, 3 women, mean age 30.5 years, range 20-49; study 3: 5 participants, 3 women, mean age 32.6 years, range 23-42) with refractory epilepsy using subdural electrode arrays implanted as part of the standard presurgical evaluation process (Supp. Table 1). Electrode array locations were thus based solely on the requirements of the clinical evaluation. Participants were recruited from three medical centers: Albany Medical Center (NY, USA), Geneva University Hospitals (Switzerland), and NYU Langone Medical Center (NY, USA). All participants gave informed consent, and the experiments reported here were approved by the respective ethical committees (Albany Medical College Institutional Review Board (Martin et al., 2016), Commission Cantonale d’Ethique de la Recherche, project number 2016-01856, and the Institutional Review Board at the New York University Langone Medical Center).

### Studies and data acquisition

Three distinct experiments were performed, one in each study center.

#### Study 1: free word repetition

The first study was a word repetition paradigm (Fig. 1a). This data appeared first in (Martin et al., 2016). The participant first heard one of six words presented through a loudspeaker (average length: 800 ms ± 20). A first cross was then displayed on the screen (1500 ms after trial onset) for 1000 ms, indicating that the participant had to imagine hearing the word. Finally, a second cross was displayed on the screen (3000 ms after trial onset) for a duration of 1500 ms, indicating that the participant had to repeat out loud the word. The six words (‘spoon’, ‘cowboys’, ‘battlefield’, ‘swimming’, ‘python’, ‘telephone’) were chosen to maximize the variability of acoustic representations, semantic categories, and number of syllables, while minimizing the variability of acoustic duration. Participants performed from 18 to 24 trials for each word.

Implanted ECoG grids (Ad-Tech Medical Corp., Racine, WI; PMT Corporation, Chanhassen, MN) were platinum-iridium electrodes (4 mm in diameter, 2.3 mm exposed) embedded in silicon. Inter-electrode distance was 4 or 10 mm. ECoG signals were recorded using seven 16-channel g.USBamp biosignal acquisition devices (g.tex, Graz, Austria) with a sampling rate of 9600 Hz. Reference and ground were chosen by selecting ECoG contacts away from epileptic foci and regions of interest. Data acquisition and synchronization with task stimuli were performed with the BCI2000 software (Schalk et al., 2004). The participant’s voice was also acquired through a dynamic microphone (Samson R21s) that was rated for voice recordings (bandwidth 80-12000 Hz, sensitivity 2.24 mV/Pa) placed 10 cm away from the patient’s face. A dedicated 16-channel g.USBamp amplifier was used to acquire and digitize the microphone signal to guarantee synchronization with ECoG data. Finally, the participants’ compliance with the imagined task was verified with an eye-tracker (Tobii T60, Tobii Sweden).

#### Study 2: rhythmic word repetition

The second study was also a word repetition paradigm (Fig. 1b). The participant first read one of twelve words presented on a laptop screen for 2000 ms. Two successive auditory cues were then presented through a loudspeaker (2100 ms and 2900 ms after the beginning of the trial). The participant then had to repeat out loud or imagine saying the word following the rhythm given by the two auditory cues (i.e. participant output was expected to start at around 3700 ms). Finally, following the same rhythm, the participant would press a key on the laptop’s keyboard (expected at around 4500 ms). Participants were repeating French words (for three participants; ‘pousser’, ‘manger’, ‘courir’, ‘pallier’, ‘penser’, ‘élire’, ‘enfant’, ‘lumière’, ‘girafe’, ‘état’, ‘mensonge’, ‘bonheur’) or similar German words (for one participant; ‘schieben’, ‘essen’, ‘laufen’, ‘leben’, ‘denken’, ‘wählen’, ‘Kind’, ‘Licht’, ‘Giraffe’, ‘Staat’, ‘Treue’, ‘Komfort’). Words were chosen to belong to four different semantic categories (concrete verbs, abstract verbs, concrete nouns, abstract nouns). Participants performed from 7 to 15 trials for each word.

ECoG signals were acquired by subdural electrode grids and strips (Ad-Tech Medical Corp; inter-electrode distance: 4 or 10 mm), amplified and digitized at 2048 Hz and stored for offline analysis (Brain Quick LTM, Micromed, S.p.A., Mogliano Veneto, Italy).

#### Study 3: rhythmic syllabic repetition

The third study was a syllable repetition paradigm (Fig. 1c). A syllable was presented rhythmically three successive times on a loudspeaker. The time interval between repetitions was selected randomly for each trial from one of three possibilities (800 ms, 1000 ms, 1200 ms). Following the same rhythm given by these syllables, the participant then had to repeat out loud or imagine saying the syllable. Participants were repeating one of three syllables (‘ba’, ‘da’, ‘ga’) in each trial. These syllables were chosen as they minimally differ acoustically (by a few dozens of ms of voice onset time, VOT) but rely on very different movements at the articulatory levels. This aims at optimizing the differences observed at the production level while limiting potential contamination by exogenous acoustic cues. Participants performed from 16 to 55 trials for each syllable.

All behavioral recordings were done via on a computer on the service tray of a hospital bed using Presentation Software (NeuroBehavioral Systems). Audio recordings were obtained using a microphone connected to the computer and were synchronized to the onset of the last auditory cue. Electroencephalographic (ECoG) activity was recorded from intracranially implanted subdural electrodes (AdTech Medical Instrument Corp.) in patients undergoing monitoring as part of treatment for pharmacologically resistant epilepsy. Electrode placement was clinically selected to localized seizure activity and eloquent tissue during stimulation mapping. Recordings included grid, depth and strip electrode arrays. Each electrode had a diameter of 4 mm (2.3 mm exposure), and the space between electrodes was 6 mm (10 mm center to center). Neural signals were recorded on a 128-channel Nicolet One EEG system with a sampling rate of 512 Hz.

### Anatomical localization of ECoG electrodes

ECoG electrodes were localized using the iELVis toolbox (http://github.com/iELVis/iELVis)(Groppe et al., 2017). Briefly, each patient’s pre-implant high-resolution structural MRI scan was automatically segmented and parcellated using Freesurfer (http://surfer.nmr.mgh.harvard.edu/)(Fischl, 2012). A post-implantation high-resolution CT or MRI scan was coregistered with the pre-implant MRI scan. Electrode artifacts were identified visually on the postimplant scan. Electrode coordinates were corrected for the brain shift caused by the implantation procedure by projecting them back to the pre-implant leptomeningeal surface. Electrode coordinates from individual participants were brought onto a common template for plotting.

### Signal processing

Time series were visually inspected, and contacts or trials containing epileptic activity and excessive noise were removed. Trials with overt speech were checked for acoustic contamination by correlating the recorded audio signal and the neural data (Roussel et al., 2020). All times series were then corrected for DC shifts by using a high-pass filter with a cutoff frequency of 0.5 Hz (zero-phase Butterworth filter of order 6, zeropole-gain design). Electromagnetic noise was removed using notch filters (forward-backward Butterworth filter of order 6, zero-pole-gain design, cutoff frequencies: 58-62 Hz, 118-122 Hz, and 178-182 Hz for studies 1 and 3; 48-52 Hz, 98-102 Hz, 148-152 Hz, and 198-202 Hz for task 2. Finally, times series were re-referenced to a common average, and downsampled to a new sampling rate of 400 Hz, 400 Hz, and 512 Hz for studies 1, 2, and 3 respectively using a finite impulse response antialiasing low-pass filter. Periods of interest for imagined and overt speech were chosen either during the period with visual cue (study 1), or 250 ms before to 250 ms after the expected production time (studies 2 and 3).

### Power spectrum

Time series were transformed to the spectral domain using an analytic Morlet wavelet transform. Power spectrum was then obtained by taking for each frequency band the average (over frequencies and time epochs of interest) of the absolute value of the complex spectral time series. We did not normalize each band independently before averaging, as normalizing caused very limited changes in the resulting powers of each band compared to when no normalization was applied. The four frequency bands of interest were the theta band (θ, 4-8 Hz), the low beta band (lβ, 12-18 Hz), the low-gamma band (lγ, 25-35 Hz), and the broadband high-frequency activity (BHA, 80-150 Hz). Cohen’s effect size 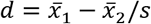 was assessed by computing the difference between the mean of the distribution of power spectrum for all trials during overt or imagined speech and the mean of the distribution of power spectrum during baseline for all corresponding trials, divided by the pooled standard deviation 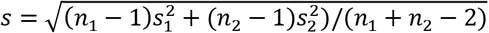, with n_*i*_ and s_*i*_ respectively the number of samples and the variance in distributions *i* ∈ {1,2}. Significance was assessed by rejecting the null-hypothesis of equality of the mean of both distributions with a two-tailed, two-sample t-test, corrected for multiple comparisons using the Benjamini-Hochberg false discovery rate (FDR) procedure (target *α* = 0.05) (Benjamini and Hochberg, 1995).

### Phase-amplitude cross-frequency coupling

Phase-amplitude cross-frequency coupling (CFC) was assessed between the phase of one band and the amplitude of another, higher-frequency band (Tort et al., 2010). To ensure that the phase of the modulating (lower) band was well defined (Aru et al., 2015), we first identified peaks in the log power spectrum for each electrode. Then, for each modulating frequency band of interest (theta band: θ, 4-8 Hz, low-beta band: lβ, 12-18 Hz, and low-gamma band: lγ, 25-35 Hz), the peak with maximal amplitude, if existing, was selected. The modulating band was then obtained by filtering original data for each modulating frequency band with a band-pass filter centered around each peak frequency with a bandwidth equal to half the size of the band of interest (i.e. 2 Hz, 3 Hz, and 5 Hz for a peak in the theta, low-beta, or low-gamma band respectively). To ensure that the modulated (higher) band was large enough to contain the side peaks produced by the modulating band, we increased the bandwidth when necessary for the modulated frequency of interest (beta band: β, 12-25 Hz, gamma band: γ, 25-50 Hz, broadband high-gamma activity: BHA, 80–150 Hz) (Aru et al., 2015). Despite those precautions, we expect that the theta/beta and low-beta/gamma phase-amplitude CFCs are not fully represented due to the limited bandwidth we can afford for the modulated frequency. The band-pass filter was a zero-phase Butterworth filter of order 6 with zero-pole-gain design. The phase and amplitude were then obtained using the Hilbert transform of the centered filtered signals.

Then, for each time epoch of interest, the histogram (18 bins) of amplitudes as a function of phases was computed and averaged across trials. Modulation index (MI) values were then calculated from the Kullback-Leibler divergence (KL) between the averaged histogram of the signal and the uniform distribution as MI = KL/log(&bins)(Tort et al., 2010). Z-scores for MI were computed by comparing the observed difference between MI values of overt/imagined time epochs and baseline *x*_*d*_ with the surrogate distribution of differences between MI values of overt/imagined time epochs and baseline *x*_*ds*_ as (*z* = *x*_*d*_ – *x*^*-*^_*ds*_)/*S*_*sd*_ with *S*_*sd*_ the standard deviation of the surrogate distribution. Surrogates were obtained by randomly shuffling 200 times the overt/imagined time epochs and baseline distribution.

One-tailed p-values corresponding to the z-scores were obtained from the cumulative normal distribution (one-tailed since the observed MI can only be greater than the surrogate one, not smaller), FDR-corrected for multiple comparisons (target *α* = 0.05) [54].

### Pairwise correlation of features with words

Pairwise correlation was quantified by computing for each speech items the Pearson correlation between power spectrum or phase-amplitude CFC features and the labels. Labels were set to 1 and -1 for the first and second word or syllable respectively of the pairwise comparison. The average pairwise correlation was then obtained for each electrode by averaging pairwise correlations across all pairs of speech items. Statistical significance was assessed by random permutations: for each pair of speech items, labels were randomly permuted, and the procedure was repeated 1000 times. A null distribution was then obtained by averaging across all speech item pairs. Significant values are those for which the p value is less than 0.05, without correction for the number of electrodes.

### Articulatory, phonetic, vocalic, and semantic representations

Words were decomposed according to their phonetic content by finding articulatory, phonetic, vocalic and semantic groups for each phoneme contained in a word (Supplementary Table 2, 3, and 4). Each word was thus represented by a set of different groups for each representation. For instance, the word ‘python’ [paɪθən] was represented as labial ([p]) and coronal ([θ], [n]) for articulatory representation, plosive ([p]), fricative ([θ]), and nasal ([n]) for phonetic representation, and low-front ([ə]), high-front ([I]), and central for the vocalic representation ([ə]). Semantic representation was only relevant for the third study, and is therefore not defined for this example. Discriminability (feature distance) between two words was then assigned to only the groups that were present in one of the two words for each representation. For instance, when comparing python ([paɪθən], that includes only labial and coronal phonemes) and cowboys ([kaʊbɔɪz], that includes only dorsal, labial, and coronal phonemes) in the articulatory representation, the feature distance was assigned to the dorsal group only, as it is the only group that discriminate both words for this representation. Discriminability to compare two words *i* and *j* was computed using the Fisher distance between their power-spectrum or cross-frequency coupling feature distributions. Fisher distance was defined as:

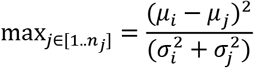

with μ_i_ and σ_i_ the mean and standard deviation of the features distribution respectively, *n*_*j*_ the dimensionality of features. Correlation could have been used as well as another metric of discriminability. The resulting values were then averaged across instances for each patient and each group. Statistical significance was assessed by random permutations: for each pair of speech items, labels were randomly permuted, and the procedure was repeated 1000 times. A null distribution was then obtained by averaging across each instance for each patient and each group. Significant values values were found after FDR-correction for multiple comparisons (target *α* = 0.05).

### Decoding

For articulatory, phonetic, and vocalic decoding, word labels were grouped together in two new classes by computing the distance between labels according to each specific representation. Distance between two words was incremented by one for each phoneme’s group that was only in one of the two words. Hierarchical clustering was then performed on the resulting distance matrix between all pairs of words (linkage criterion that uses the maximum distances between all observations of the two sets of observations). The new classes were selected by taking groups of words that were close-by in the dendrogram, while minimizing the class imbalance. For semantic decoding, words labels were grouped into two classes, following the initial experimental design. The ‘abstract’ class contains the words: ‘pousser’, ‘manger’, ‘courir’, ‘enfant’, ‘lumière’, ‘girafe’. The ‘concrete’ class contains the words: ‘pallier’, ‘penser’, ‘élire’, ‘état’, ‘mensonge’, ‘bonheur’.

For each binary classification problem resulting of this clustering procedure, we trained a classifier. We used a 10-fold cross-validation approach, i.e. data was divided in 10 blocks, with 90% of the blocks being used for training, and the remaining block being used for testing. This procedure was repeated 10 times by shifting every time the block used for testing. We used a support vector machine algorithm with a linear kernel for classification. Feature selection was done using recursive feature elimination, (starting with the full set of features and removing sequentially features that do not contribute to the classifier performance). Feature selection was done using nested 5-fold cross-validation within the training set. Score was evaluated using balanced accuracy to account for class imbalance that could occur when there were more samples in one of the two classes.

Thresholds for significant classification performance were obtained independently for each subject from an inverse binomial distribution, which accounts for the possibility of obtaining by chance accuracies higher that 50% in a binary classification problem because of a low number of trials (Combrisson and Jerbi, 2015).

### Code and data availability

Code was written in MATLAB and Python, and is available at (#URL will be made available upon publication). Ethical and privacy imperatives prevent us from posting patient-related data to public repositories. Requests for data should be directed to Dr. Mégevand.

