## Supplementary Material for "Imagined speech can be decoded from low- and cross-frequency features in perceptual space"

Timothée Proix, Jaime Delgado, Andy Christen, Stephanie Martin,
Xing Tian, David Poeppel, Werner K. Doyle, Orrin Devinsky,
Luc Arnal, Pierre Mégevand, Anne-Lise Giraud

#### Realigning imagined speech trials

Because of the absence of any behavioral output during imagined speech, combined with a very low signal-to-noise ratio, realigning imagined speech trials is a challenging task; previous attempts have used for instance dynamic time warping approach between each pairs of trials to build similarity matrix across trials (Martin et al., 2016, 2014). We have tried several approaches, using for instance dynamic time warping variations to directly average across trials (Morel et al., 2018), or simple power comparisons, without much success (results not shown).

### Supplementary figures

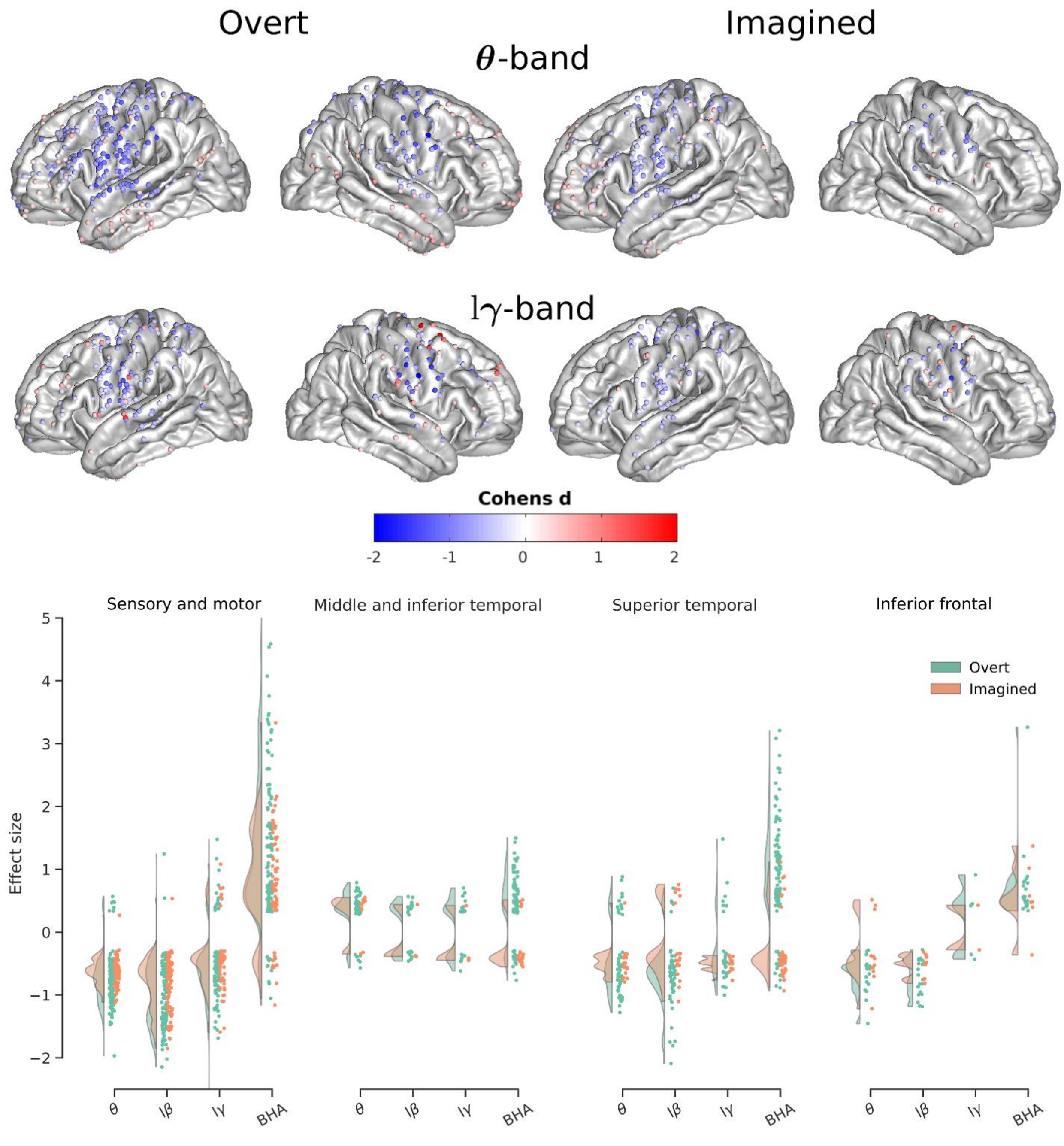


Figure 1: **Power spectrum changes during overt and imagined speech compared to baseline.** Effect sizes for each ROI and frequency band.

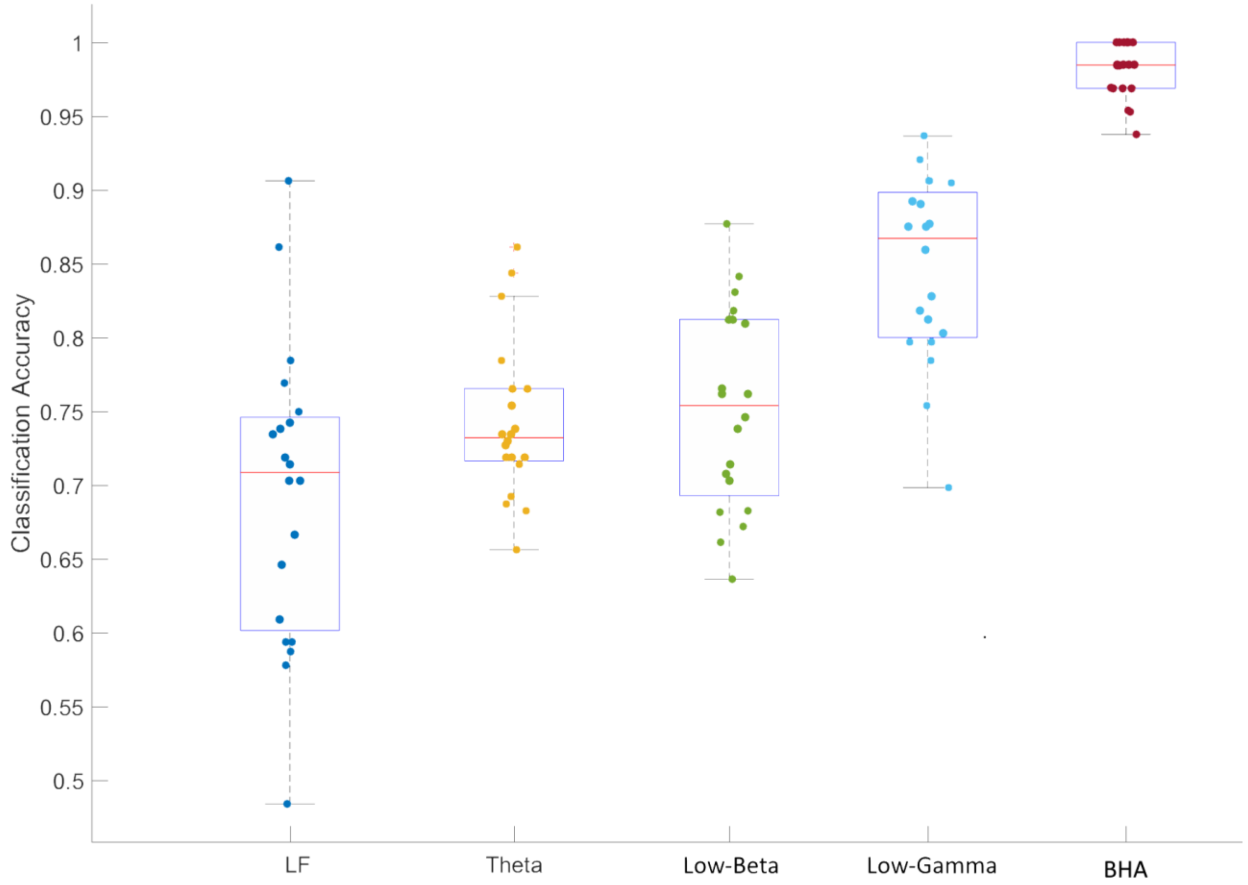


Figure 2: **Discriminating among tasks (listening vs overt vs imagined) using classification with power features.** Features correspond to concatenated time series for power in the different frequency bands. Low-frequency (LF) features correspond to ECoG signal amplitude filtered below 20 Hz using a zero-phase Butterworth filter of order 8, rather than power. A linear discriminant analysis classifier was trained using using a 5-fold cross-validation.

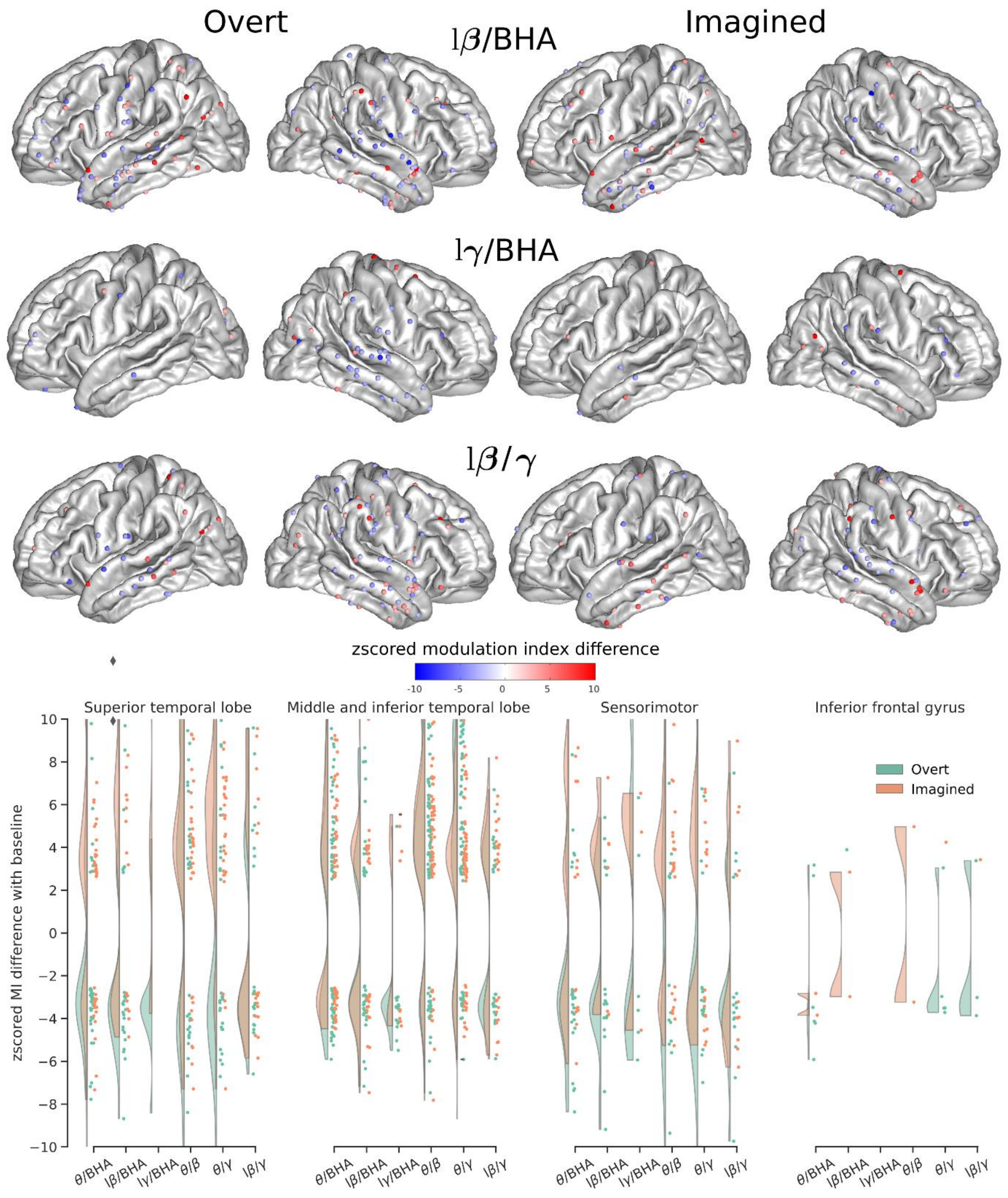


Figure 3: **Cross-frequency coupling between the phase of one frequency band and the amplitude of another frequency band in the same contact** Zscored modulation index for each ROI and frequency band

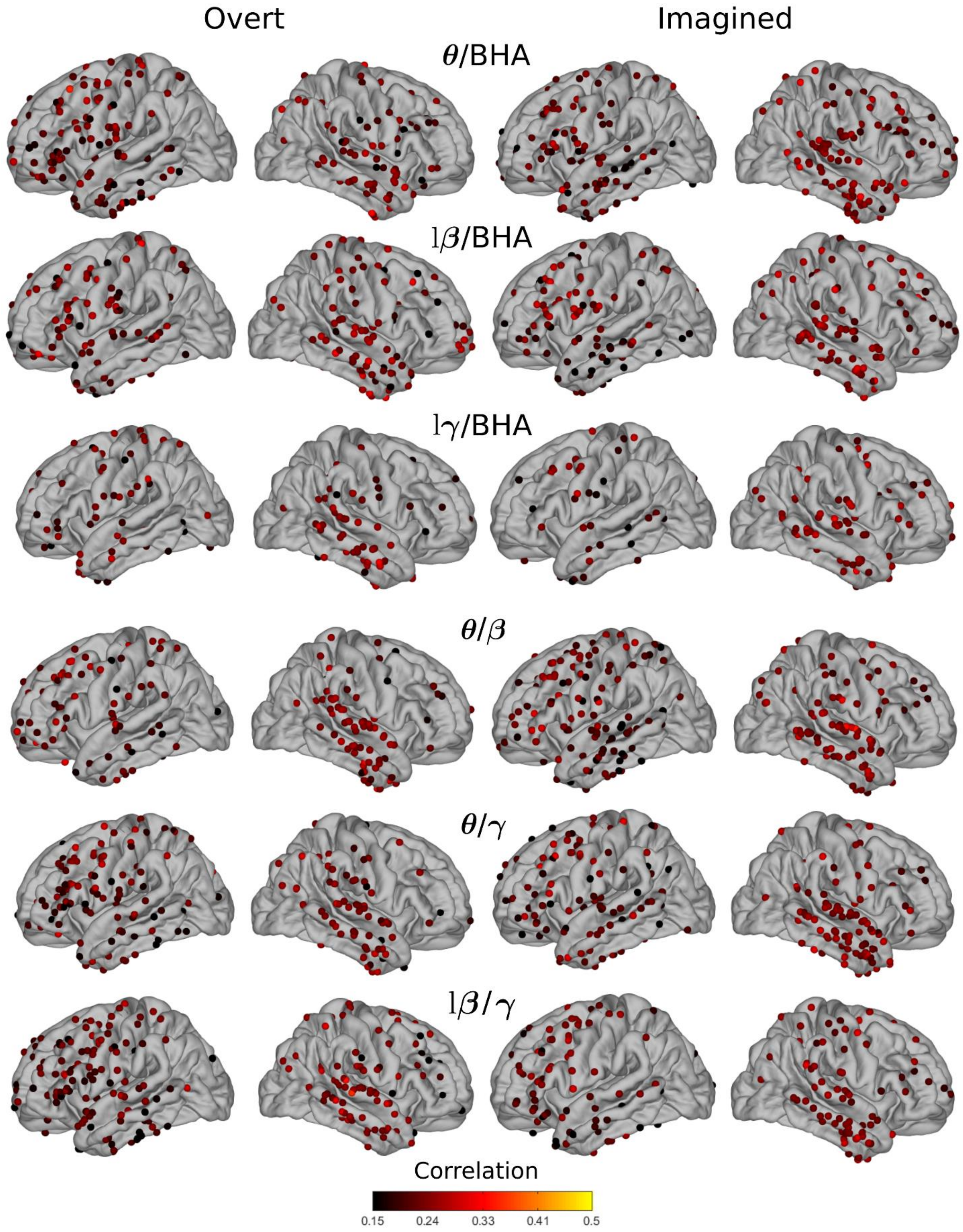


Figure 4: **Average correlations between speech and phase-amplitude coupling features.** Pairwise correlations between words/syllables and phase-amplitude coupling features averaged across all pairs for overt and imagined speech.

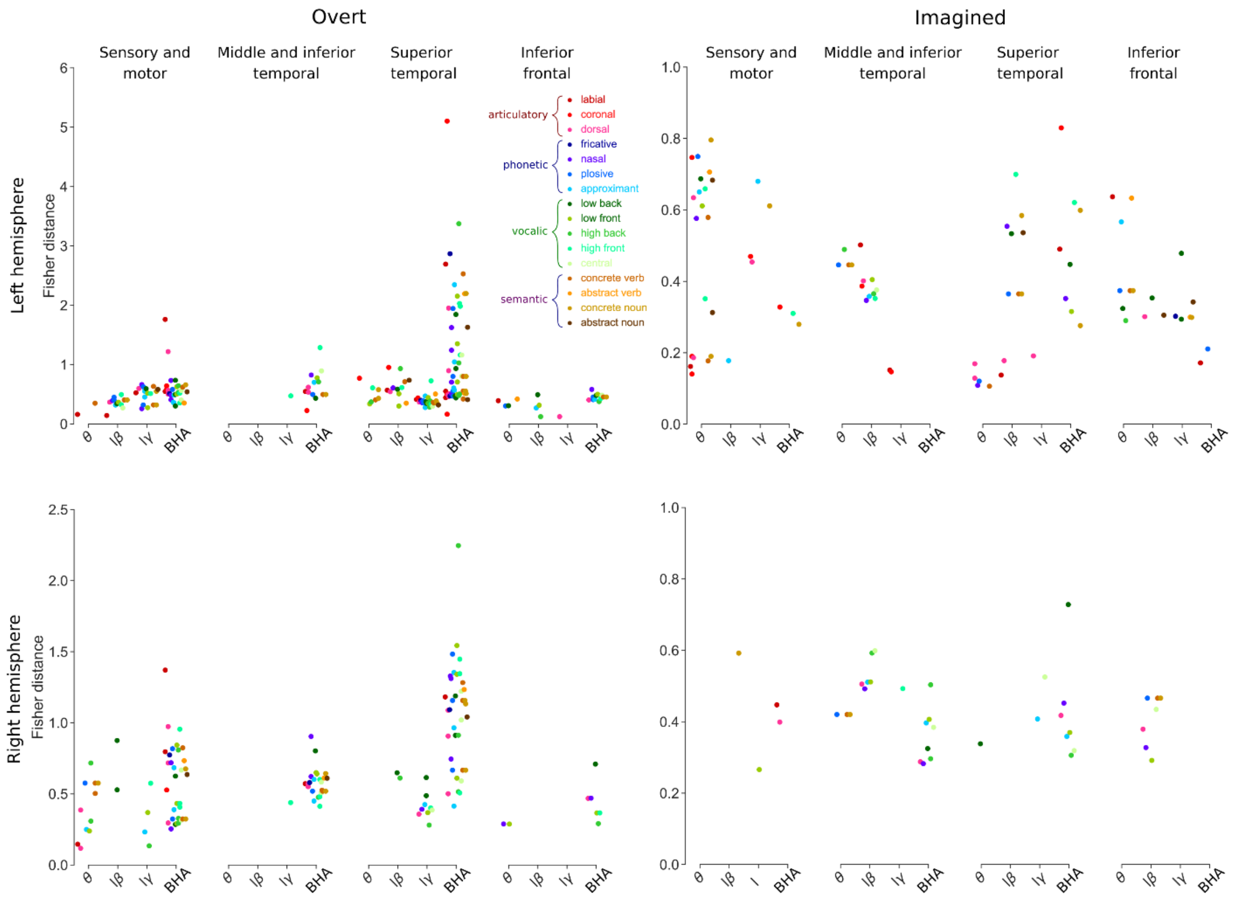


Figure 5: **Separation of power spectrum changes for articulatory, phonetic and vowel representations in different brain regions and frequency bands.** Only significant values are shown (permutation test, FDR corrected, target threshold $\alpha = 0.05$). Note the different scales between overt and imagined speech.

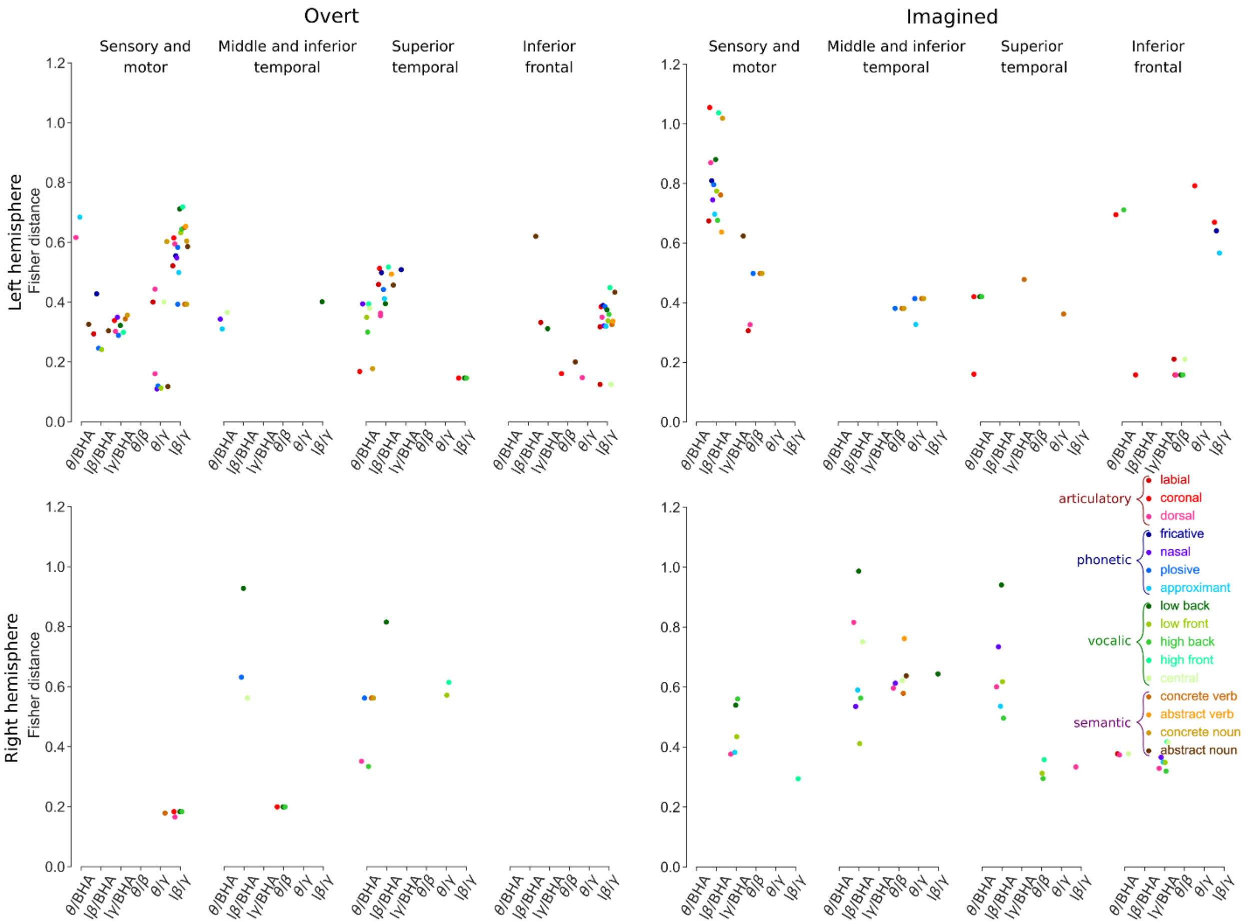


Figure 6: **Separation of power spectrum changes for articulatory, phonetic and vowel representations in different brain regions and frequency bands.** Only significant values are shown (permutation test, FDR corrected, target threshold $\alpha = 0.05$).

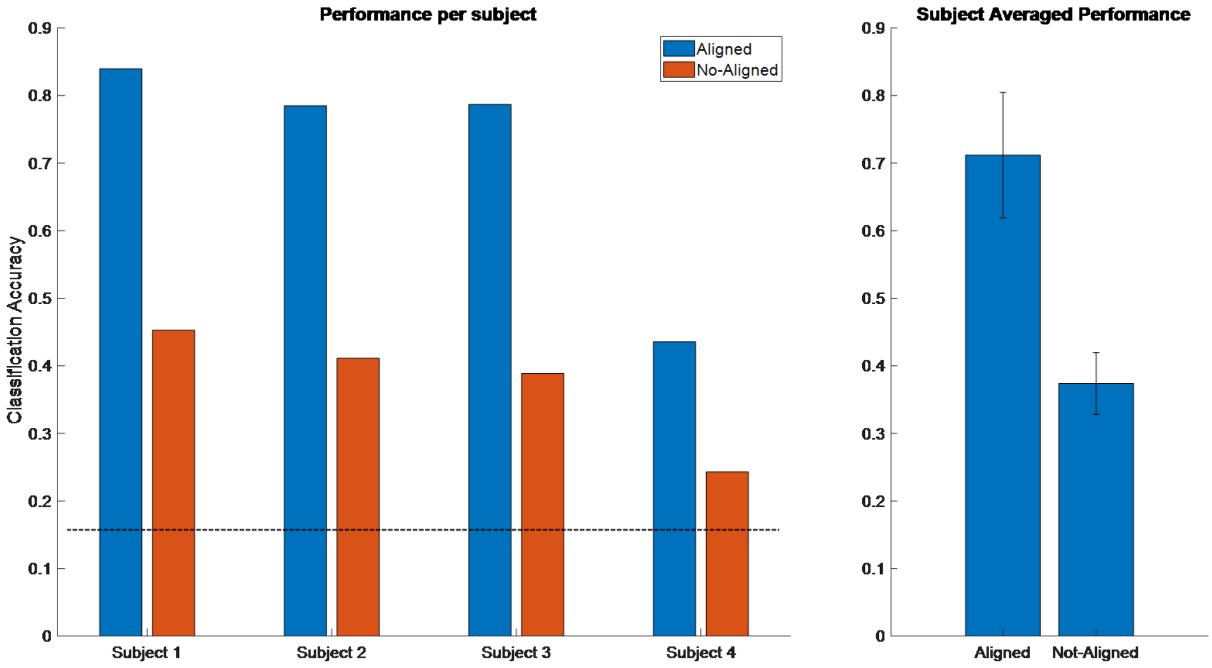


Figure 7: **Decoding performance of overt speech increases significantly after realigning trials for study 1.** Multiclass classification results (6 classes). Trials were aligned on the onset of audio recordings during production. A linear discriminant function was trained using the electrodes that show activity during the task with respect to baseline. Classification performance was obtained using a stratified 10-fold cross-validation, using each time 90% of the data for training and 10% for testing.

| Study | Patient | Age | Sex | Handedness | Speech lateralization |
|---|---|---|---|---|---|
| Study1 | P1 | NA | F | L | L (iEEG) |
| | P2 | 25 | F | R | R (iEEG) |
| | P3 | 19 | F | L | L (iEEG) |
| | P4 | 33 | F | R | R (iEEG) |
| Study2 | P5 | 22 | M | R | L (fMRI) |
| | P6 | 49 | F | L | L (fMRI) |
| | P7 | 30 | F | R | L (fMRI, ESM) |
| | P8 | 20 | F | L | bilateral , R-dominant (fMRI), not L-dominant (ESM) |
| Study3 | P9 | 23 | F | R | bilateral |
| | P10 | 42 | M | R | unknown |
| | P11 | 42 | F | R | L |
| | P12 | 31 | M | R | L |
| | P13 | 25 | F | R | L |

Table 1: **Clinical information for participants in the three studies.** L: left, R: right.

| | **Phoneme** | | | | | | | | | | | | | | | | | | | | | |
|---|---|---|---|---|---|---|---|---|---|---|---|---|---|---|---|---|---|---|---|---|---|---|
| | b | d | f | g | k | l | m | n | p | K | s | t | v | z | T | N | j | z | S | ¸c | r | w |
| **Articulatory** | | | | | | | | | | | | | | | | | | | | | | |
| Labial | x | | x | | | | x | | x | | | | x | | | | | | | | | |
| Coronal | | x | | | | x | | x | | | x | x | | x | x | | | x | x | | x | |
| Dorsal | | | | x | x | | | | | x | | | | | | x | x | | | x | | x |
| **Phonetic** | | | | | | | | | | | | | | | | | | | | | | |
| Nasal | | | | | | | x | x | | | | | | | | x | | | | | | |
| Plosive | x | x | | x | x | | | | x | | | x | | | | | | | | | | |
| Approximant | | | | | | x | | | | | | | | | | | x | | | | | x |

Table 2: **Articulatory and phonetic representation for each phoneme used in the tasks.**

| | **Phoneme** | | | | | | | | | | | | | | |
|---|---|---|---|---|---|---|---|---|---|---|---|---|---|---|---|
| | i | y | I | Y | e | E | æ | œ | a | @ | u | U | O | ˜O | A˜ |
| **Vocalic** | | | | | | | | | | | | | | | |
| Front | x | x | x | x | x | x | x | x | x | | | | | | |
| Middle | | | | | | | | | | x | | | | | |
| Back | | | | | | | | | | | x | x | x | x | x |
| High | x | x | x | x | x | | | | | | x | x | | | |
| Central | | | | | | | | | | x | | | | | |
| Low | | | | | | x | x | x | x | | | | x | x | x |

Table 3: **Vocalic representation for each vowel used in the tasks.**

| | **Word** | |
|---|---|---|
| | Concrete | Abstract |
| Verb | pousser, manger, courir | pallier, penser, élire |
| Noun | enfant, lumière , girafe | état, mensonge, bonheur |

Table 4: **Semantic representation for words used in study 2.**
